# Building applications for interactive data exploration in systems biology

**DOI:** 10.1101/141630

**Authors:** Bjørn Fjukstad, Vanessa Dumeaux, Karina Standahl Olsen, Michael Hallet, Eiliv Lund, Lars Ailo Bongo

**Affiliations:** Department of Computer Science, UiT The Arctic University of Norway; Department of Biology, Concordia University; Department of Community Medicine, UiT The Arctic University of Norway

## Abstract

As the systems biology community generates and collects data at an unprecedented rate, there is a growing need for interactive data exploration tools to explore the datasets. These tools need to combine advanced statistical analyses, relevant knowledge from biological databases, and interactive visualizations in an application with clear user interfaces. To answer specific research questions tools must provide specialized user interfaces and visualizations. While these are application-specific, the underlying components of a data analysis tool can be shared and reused later. Application developers can therefore compose applications of reusable services rather than implementing a single monolithic application from the ground up for each project.

Our approach for developing data exploration applications in systems biology builds on the microservice architecture. Microservice architectures separates an application into smaller components that communicate using language-agnostic protocols. We show that this design is suitable in bioinformatics applications where applications often use different tools, written in different languages, by different research groups. Packaging each service in a software container enables re-use and sharing of key components between applications, reducing development, deployment, and maintenance time.

We demonstrate the viability of our approach through a web application, MIxT blood-tumor, for exploring and comparing transcriptional profiles from blood and tumor samples in breast cancer patients. The application integrates advanced statistical software, up-to-date information from biological databases, and modern data visualization libraries.

The web application for exploring transcriptional profiles, MIxT, is online at mixt-blood-tumor.bci.mcgill.ca and open-sourced at github.com/fjukstad/mixt. Packages to build the supporting microservices are open-sourced as a part of Kvik at github.com/fjukstad/kvik.

## Introduction

In recent years the biological community has generated an unprecedented ammount of data. While the cost of data collection has drastically decreased, data analysis continue to be a larger fraction of the total experiment cost.[1] An important part of data analysis includes the time spent by human experts interpreting the results. This calls for novel methods for building data analysis tools for data exploration and interpretation.

In the field of systems biology, data exploration applications need to link results to relevant prior knowledge. In the later years there has been a tremendous effort to curate databases with relevant information on genes and processes. Databases such as the Molecular Signatures Database (MSigDB)^1^ and the Kyoto Encyclopedia of Genes and Genomes (KEGG)^2^ both provide interfaces to retrieve data that can be used to better the understanding of data analysis results.

Several tools for biological data analysis are now available in various programming languages. These include a wide variety of bioinformatics methodologies and graphical analysis tools. In the R statistical programming language developers share software through repositories such as CRAN^3^ or Bioconductor^4^. In other languages, libraries for biological bom-putation are often availalbe like BioPython[2] and biogo[3] for Python or Go, respectively. Projects such as Galaxy^5^ and Common Workflow Language (CWL)^6^ enable resarchers to build and run biological data analysis pipelines consisting of a wide range of tools. Although these framework are tremendously helpful, we need novel approaches to build applications that integrate high-performance bioinformatics tools with specialized user-interfaces and interactive visualizations.

Different programming languages solve different tasks. For example, new biological data analysis techniques are quickly realeased in R and its package repositories; high-performance computer vision tasks are performed in C++ and OpenCV; and portable user interfaces more easily built in HTML, CSS and JavaScript. Therefore applications that integrate novel statistical analysis tools, interactive visualizations, and biological databases likely need to include several components written in different languages.

A microservice architecture structures an application into small reusable, loosely coupled parts. These communicate via lightweight programming language-agnostic protocols such as HTTP, making it possible to write single applications in multiple programming languages. This way the most suitable programming language is used for each specific part. To build a microservice application, developers bundle each service in a container. Containers are built from configuration files which describe the operating system, software packages and versions of these. This makes reproducing the analyses, database lookups, library versions in an application a trivial task. The most popular implementation of a software container is Docker^7^, but others such as Rkt^8^ exist. Initiatives such as BioContainers^9^ now provide containers pre-installed with different bioinformatics tools. While the enabling technology is available, the microservices approach is not yet widely adopted in bioinformatics.[4]

From our experience we have identified a set of components and features we needed to build data exploration applications.

1. A low-latency language-independent approach for integrating, or embedding, statistical software, such as R, directly in a data exploration application.
2. Low latency language-independent interface to online reference databases in biology that users can query to better understand resulst from statistical analyses.
3. A simple method for deploying and sharing the components of an application between projects.

In this paper, we describe a novel approach for building data exploration applications in systems biology. We show that by building applications as a set of services we can reuse and share its components between applications. The key services of a biological data exploration application are i) a compute service for executing statistical analyses in languages such as R, ii) a database query service for retrieving information from biological databases, and iii) the user-facing visualizations and user-interfaces. In addition, by packaging the services using container technology they are easy to deploy, simple to reproduce, and easy to share between projects. We have used our approach to build a number of applications, both command-line and web-based. In this paper we describe how we used our approach to develop MIxT, a web application for exploring and comparing transcriptional profiles from blood and tumor samples.

## Methods

In this section we first motivate our microservice approach based on our experiences developing the MIxT web application. We describe the process from initial data analysis to the final application, highlighting the importance of language-agnostic services to facilitate the use of different tools in different parts of the application. We show that these services are easy to deploy and provide performance to use them to build data exploration applications. We then generalize the ideas to a set of principles and services that can be reused and shared between applications, and show their design and implementation.

### Motivating Example

The aim of the Matched Interactions Across Tissues (MIxT) study was to identify genes and pathways in the primary breast tumor that are tightly linked to genes and pathways in the patient blood cells.[5] We generated and analyzed expression profiles from blood and matched tumor cells in 173 breast cancer patients included in the Norwegian Women and Cancer (NOWAC) study. The MIxT analysis starts by identifying sets of genes tightly co-expressed across all patients in each tissue. Each group of genes or modules were annotated based on known a priori biological knowledge about gene functionality. Focus was placed on the relationships between tissues by asking if specific biologies in one tissue are linked with (possibly distinct) biologies in the second tissue, and this within different subgroup of patients (i.e. subtypes of breast cancer).

We built an R package^10^ with the statistical methods and static visualizations for identifying associations between modules across tissues. The exploration of the results encompass the examination of ~ 20 modules and their functional enrichments. That is ~ 23 × 19 = 437 associations computed for each of the 22 patient subgroups.

To explore this large amount of data and results we needed an interactive point-and-click application that could interface with the statistical methods and link the results to online databases. This would make it possible for users to explore the results without any coding background. We needed an application that could interface with MSigDB to fetch gene set metadata and the Entrez Programming Utilities (E-utils) to retrieve gene meta-data. To make the interfaces to the statistical analyses and databases re-usable by other applications that we aim to implement later it was necessary for these to communicate using open protocols.

In addition, we required that the application was containerized. This allows us to deploy the application on a wide range of hardware, from local installations to deployments to cloud providers such as Amazon Web Services (AWS)^11^.

### Design

Our experience can be generalized into the following design principles for building applications in bioinformatics:

Principle 1: Build applications as collections of language-agnostic microservices. This enables re-use of components and does not enforce any specific programming language on the user-facing logic or the underlying components of the application. Within bioinformatics, researchers and application developers use a wide range of programming languages to build tools. By composing an application of services communicating over standard protocols it is possible to re-use existing tools in new applications.
Principle 2: Use software containers to package each service. This has a number of benefits; it simplifies deployment, ensures that dependencies and libraries are installed, and it simplifies sharing of services between developers. Using these design principles we built the MIxT web application using the microservices in Kvik to interface with statistical analyses and biological databases. Kvik provides a compute service for executing statistical analyses and a database service for retrieving relevant information on genes and biological processes. Using these it is possible to develop specialized data exploration application in any modern programming language. In the rest of the section we discuss the design choices we made to build the microservices that power the MIxT web application.

### Compute Service

The main cornerstone of every data exploration application in systems biology is a dataset. Datasets go through a series of transformations before they can be interpreted by experts. Because of the complexity of these datasets, package repositories such as BioConductor provide software for reading and analyzing the data. Because of the complex datasets, there are a number of requirements for systems that can explore them i) datasets require specialized software to be read, ii) datasets can be too big to fit on a desktop computer, iii) statistical methods for analyzing the datasets are too computationally intensive to run on a desktop computer, iv) users may want to modify statistical analyses directly from an application, and v) users need to know exactly what transformations has been done to a dataset to reproduce the analyses.

From these requirements we have designed the compute service in Kvik. The compute service interfaces directly with the R programming language, making it possible to call functions from any R package. Application developers can use the compute service to execute analyses and return results. By interfacing directly with R developers can leverage this to produce dynamic applications. For example, if an application uses a clustering method to color nodes in a graph, end-users can tweak parameters that interactively within the application that changes the node coloring in graph visualization. Also, by interfacing with R directly from an application, the application can store provenance data on the statistical methods and input parameters being used. By placing data storage and analysis into a service, application developers can deploy it on a powerful server while keeping the user-facing application logic on the desktop. Packaging the service into a software container also provides the necessary functionality to ensure that the statistical analyses can be reproduced later.

The compute service offers three main operations to interface with R. i) to call a function from an R package, ii) to get the results from a previous function call, and iii) a catch-all term that both calls a function and returns the results. We use the same terminology as OpenCPU[6] and have named the three operations Call, Get, and RPC respectively. These three operations provide the necessary interface for applications to include data in the applications.

### Database Service

To understand analysis results experts query databases and scientific literature. There are a wealth of online databases, some of which provide open APIs in addition to web user interfaces that application developers can make use of. While the databases can provide helpful information, there are some challenges including them in interactive data exploration applications:

i. the APIs aren’t fast enough to use in interactive applications where the application has to perform multiple database calls,
ii. some databases put restrictions on the number of database calls, and
iii. there is no uniform way for storing database lookup provenance to reproduce the database lookups.

To solve these problems we built a database service for application developers to include. The service includes a simple caching mechanism that solves the two challenges. If we cache queries to the database we can speed up subsequent calls and reduce the load on the respective databases. Both the query from the application and the response from the databases are stored for later use. The database service provides an open HTTP interface to biological databases for retrieving meta-data on genes and processes. We have currently packages for interfacing with E-utilities^12^, MSigDB^13^, Hugo Gene Nomenclature Committe (HGNC)^14^, and Kyoto Encyclopedia of Genes and Genomes (KEGG)^15^.

### Applications

Kvik provides services to perform database lookup and execute statistical analyses. Since we provide both services as software containers, application developers simply run these and write applications that interface with them. Using our services application developers have a platform for running statistical analyses, and a fast clean interface to get metadata about genes and processes.

Figure 1 shows how the the MIxT application is built using Kvik microservices. In MIxT we built a specialized web application that interfaces with Kvik to get data from biological databases and to run statistical analyses from the mixt R package^16^.

**Figure 1:**
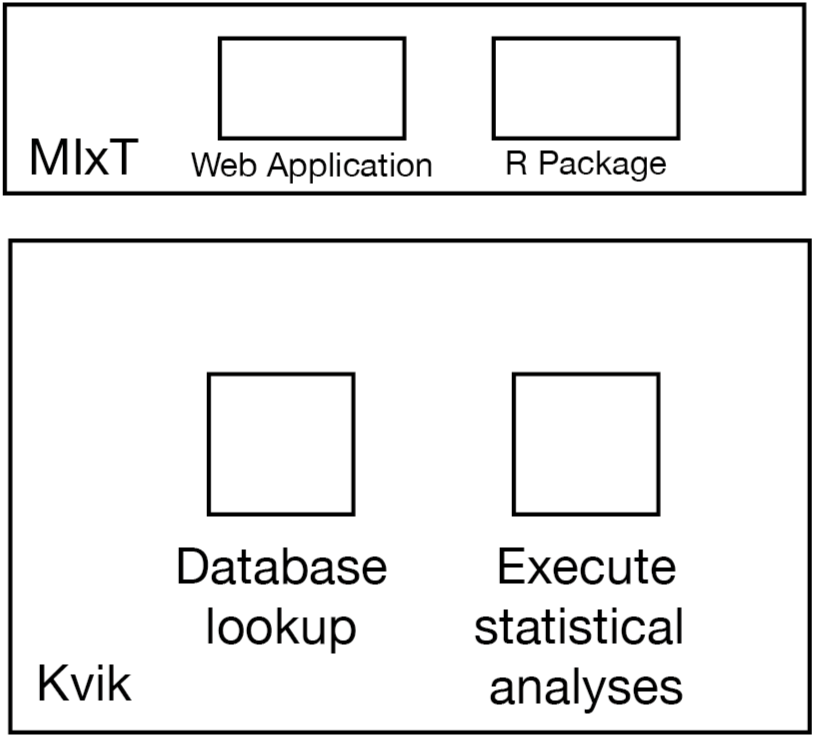
An overview of the relationship between the MIxT application and Kvik. MIxT contains a web application (online at mixt-blood-tumor.bci.mcgill.ca) and the R Package that provides analyses and data to the web application. Kvik provides the services for running the statistical analyses from the R package, and the database lookups found in the web application.

### Implementation

In ths section we describe the implementation details of the microservices we provide in Kvik.

Kvik is implemented as a collection of Go packages required to build services that can integrate statistical software in a data exploration and provide an interface to up-to-date biological databases. We chose the Go programming language because of its performance, ease of development, and simple deployment. To integrate R we provide two packages *gopencpu* and *r*, that interface with OpenCPU and Kvik R servers respectively. To interface with biological databases we provide the packages *eutils, gsea, genenames,* and *kegg* that interface with E-utils, MsigDB, HGNC and KEGG respectively. In addition to these packages we provide Docker images that implement the two required microservices.

Both the compute and the databases service in Kvik builds on the standard *http* package in Go. The database service use the *gocache*^17^ package to cache any query to an online database. In addition we deploy each service as Docker containers.^18^.

### Compute Service

The compute service is an HTTP server that communicates with a pre-set number of R processes to execute statistical analyses. On start of the compute service launches a user-defined number of R worker sessions for executing analyses, default is 5. The compute service uses a round-robin scheduling scheme to distribute incoming requests to the workers. We provide a simple FIFO queue for queuing of requests. The compute service also provides the opportunity for users to cache analysis results to speed up subsequent calls.

The compute service in Kvik is built using a hybrid state pattern. A hybrid state pattern origins from functional programming, where output from a method only depends on its inputs and not the program state.[6]. In practice this means that the compute services stores the output results from function calls, but can compute them if the service has to restart. This makes it possible for us to scale the compute service horizontally to handle more requests. If an R worker session for some reason crashes the compute service simply starts up a replacement.

## Matched interactions across tissues

We show the viability of the microservices approach in Kvik by describing the MIxT web application for exploring and comparing transcriptional profiles from blood and tumor samples. We also evaluate the performance of the microservices in Kvik to show its usefulness in building MIxT.

### Analysis Tasks

The web application provides functionality to perform six data analysis tasks (A1-A6):

**A1:** Explore co-expression gene sets in tumor and blood tissue. We simplify the process of exploring the computed co-expression gene sets, or modules, through the web-application. The application visualize gene expression patterns together with clinico-pathological variables for each module. In addition we enable users to study the underlying biological functions of each module by including gene set analyses between the module genes and known gene sets.
**A2:** Explore co-expression relationships between genes. Users can explore the co-expression relationship as a graph visualization. The network visualizes each gene as a node and a significant co-expression relationship as an edge.
**A3:** Explore relationships between modules from each tissue. Users can explore the relationship between modules from different tissues. We provide two different metrics to compare modules, and the web application enables users to interactively browse these relationships. In addition to providing visualizations the compare modules from each tissue, users can explore the relationships, but for different breast cancer patient groups.
**A4:** Explore relationships between clinical variables and modules. In addition to comparing the association between modules from both tissues, users also have the possibility to explore the association with a module and a specific clinical variable. It is also possible to explore the associations stratifying on breast cancer patient group.
**A5:** Explore association between user-submitted gene lists and computed modules. We want to enable users to explore their own gene lists to explore them in context of the co-expression gene sets. The web application must handle uploads of gene lists and compute association between the genelist and the MIxT modules on demand.
**A6:** Search for genes or gene lists of interest. To facilitate faster lookup of genes and biological processes, the web application provides a search functionality that lets users locate genes or gene lists and show association to the co-expression gene sets.

### Design and Implementation

From these six analysis tasks we designed and implemented MIxT as a web application that integrates statistical analyses and information from biological databases together with interactive visualizations. The MIxT web application consists of three services: i) the web application itself containing the user-interface and visualizations; ii) the compute service performing the MIxT analyses delivering data to the web application; and iii) the database service providing up-to-date information from biological databases. Each of these services run within Docker containers making the process of deploying the application simple.

We structured the MIxT application with a separate view for each analysis task. To explore the coexpression gene sets (**A1**) we built a view that combines both static visualizations from R together with interactive tables with gene overlap analyses. Figure 3 shows the web page presented to users when they access the co-expression gene set ‘darkturquoise’ from blood. Using the Kvik compute service we can generate plots on demand and provide users with highresolution PDFs or PNG files. To explore the coexpression relationship between genes (**A2**) we use an interactive graph visualization build with Sigmajs^19^. We have built visualization for both tissues, with graph sizes of 2705 nodes and 90 348 edges for the blood network, and 2066 nodes and 50 563 edges for the biopsy network. The sigmajs visualization library has functionality for generating a layout for large networks, but we generate this layout server-side to reduce the computational load on the client. To generate this layout we use the GGally package^20^. By generating the network layout using the compute service we relieve the clients.

To visualize relationships between modules from different tissues (**A3**), or their relationship to clinical variables (**A4**) we built a heatmap visualization using the d3^21^ library. Figure 2 shows an example of this heatmap visualization, showing the association between the clinical variables and the modules from biopsy for all samples.

**Figure 2:**
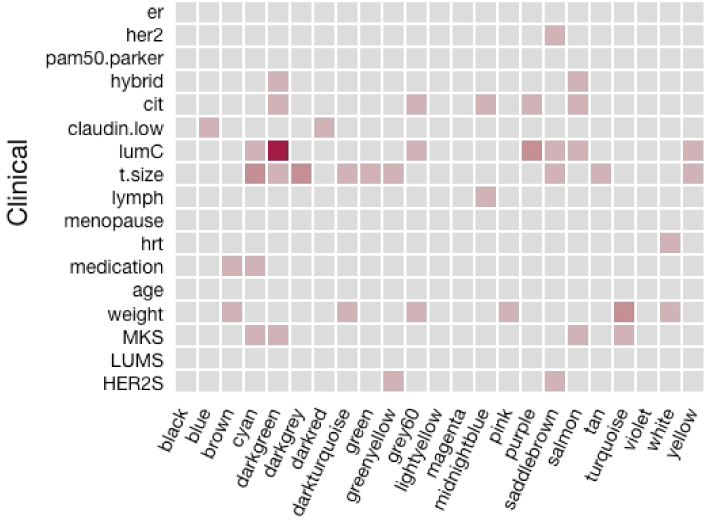
Heatmap visualization of the association between clinical variables and the modules in biosy. The visualization is built using the d3 JavaScript library. The visualization can be viewed online at mixt-blood-tumor.bci.mcgill.ca/clinical-comparison

Since we interface directly with R we can run analyses on demand. We built a simple upload page where users can upload their gene sets or type them in manually (**A5**). The file is uploaded to the web application which redirects it to the compute service that runs the analyses. Similarly we can take user input to search for genes and processes (**A5**).

### Evaluation

To investigate if it is feasible to implement parts of an application as separate services, we evaluate the response times for a set of queries. We have also investigated the time the MIxT web application use to produce the visualizations for the analysis tasks.

To evaluate the database service we measure the query time for retrieving information about a specific gene with and without caching. This illustrates how we can improve performance in an application by using a database service rather than accessing the database directly. We evaluate the query time for 1, 2, 5, 10, and 15 concurrent requests. Since the database service is just a lightweight HTTP server we use a AWS EC2 *t2.micro*^22^ instance to host it.

From the results in Table 1 we see a significant improvement in response time when the database service caches the results from the database lookups. In addition by serving the results out of cache we reduce the number of queries to the online database down to one.

**Table 1:**
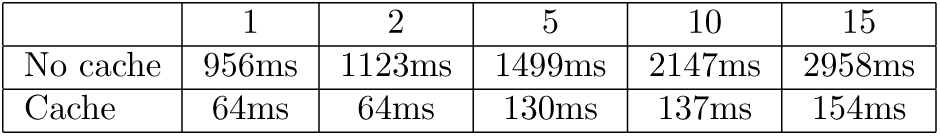
Time to retrieve a gene summary for a single gene, comparing different number of concurrent requests.

|  | 1 | 2 | 5 | 10 | 15 |
| --- | --- | --- | --- | --- | --- |
| No cache | 956ms | 1123ms | 1499ms | 2147ms | 2958ms |
| Cache | 64ms | 64ms | 130ms | 137ms | 154ms |

We evaluate the compute service by running a small microbenchmark. The benchmark consists of two operations: first generate a set of numbers, then plot them and return the resulting visualization. We show that the latency is low enough to use it in an interactive application, and compare our solution to the OpenCPU system. To show that it performs well under heavy load we perform the same operations using 1, 2, 5, 10, and 15 concurrent requests. We use two *c4.large* instances on AWS EC2 running the Kvik compute service and OpenCPU base docker containers. The servers have caching disabled.

With single requests the the mean execution time in Kvik is 274ms while OpenCPU uses 500ms. We then investigate how each service handles concurrent requests. Table 2 shows the time to complete the benchmark for different number of concurrent connections. We see that the compute service in Kvik performs better than the OpenCPU alternative.

**Table 2:**
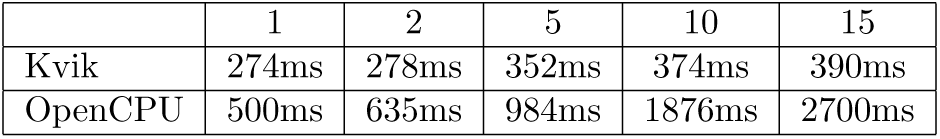
Time to complete the R microbenchmark with different number of concurrent connections.

We also investigate the performance of the MIxT web application to discover potential areas of improvement. We measured the time from an user clicks on a link to open a specific view, until the user can interact with the results. Table 3 show the results from our evaluation, with anlysis tasks A1 and A2 being the most time consuming. A1 generates the view in Figure 3 which contains large HTML tables with results from the gene set tests that take the major fraction of the time to completion. The time to completion in analysis task A2 comes from retrieving and rendering the two large graphs.

**Figure 3:**
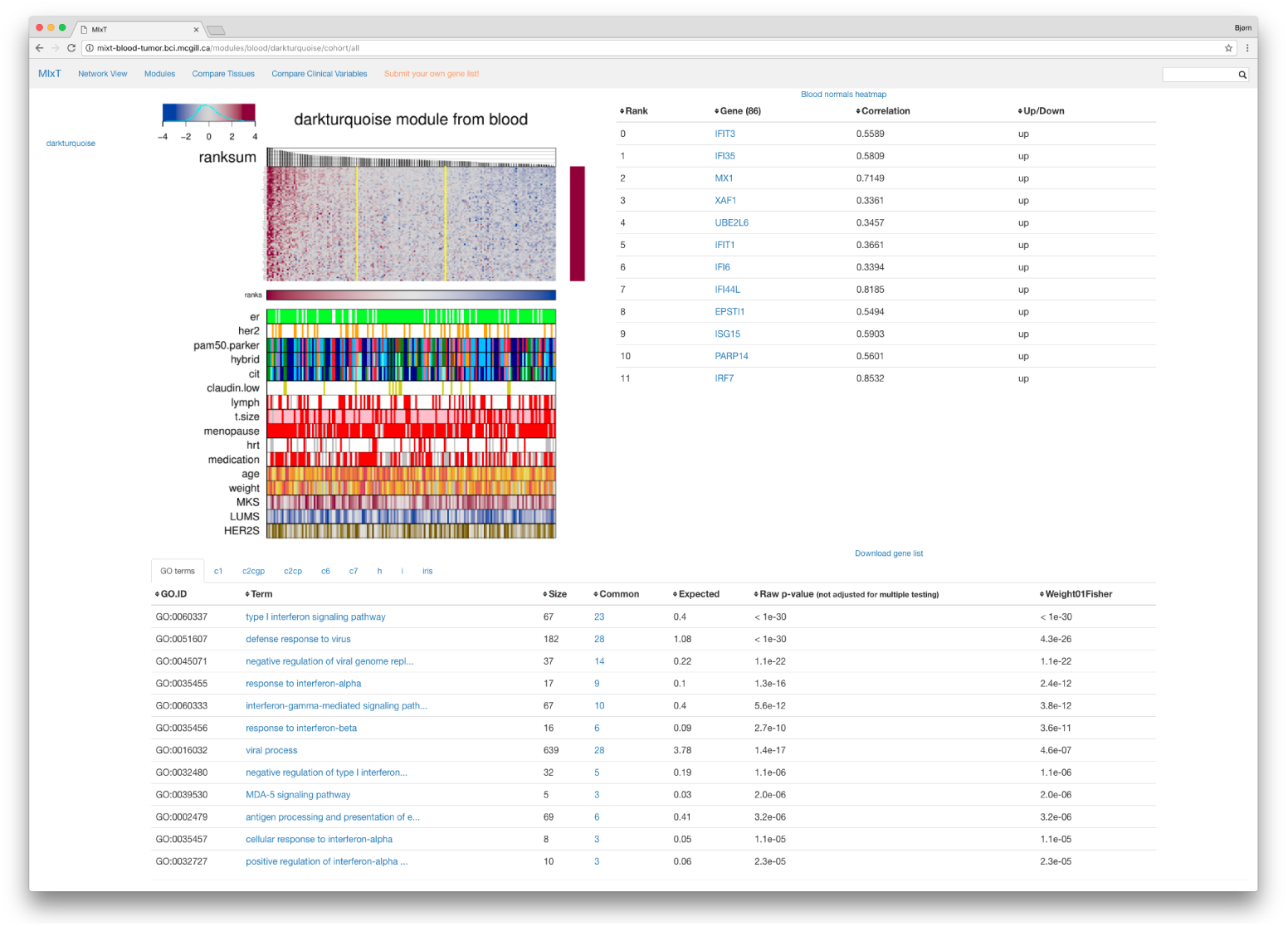
MIxT module overview page. The screenshot show the user interface for exploring a single module. It consists of three panels. The top left panel contains the gene expression heatmap. The top right panel contains a table of the genes found in the module. The bottom panel contains the results of gene overlap analyses from the module genes and known gene sets from MSigDB. The visualization is online at mixt-blood-tumor.bci.mcgill.ca/modules/blood/darkturquoise/cohort/all

**Table 3:** An overview of the number of calls to the compute service and completion time before the results for the different analysis tasks are ready to be explored. The number of calls and completion time for analysis task 6 depends on the search query.

| Analysis task | Number of calls | Time to completion |
| --- | --- | --- |
| A1 | 10 | 10 seconds |
| A2 | 5 | 30 seconds |
| A3 | 1 | 2 seconds |
| A4 | 1 | 2 seconds |
| A5 | 2 | 3 seconds |
| A6 | NA | NA |

## Related Work

In this section we discuss methods for integrating statistical analyses in data exploration applications, different visualization frameworks, interfaces to biological reference databases, and methods for containerizing applications.

### Integrate Statistical Analyses

Shiny is a web application framework for R^23^ It allows developers to build web applications in R without having to have any knowledge about HTML, CSS or Javascript. Its widget library to provides more advanced Javascript visualizations such as Leaflet^24^ for maps or three.js^25^ for WebGL-accellerated graphics. Developers can choose to host their own web server with the user-built Shiny Apps, or host them on public servers. Shiny forces users to implement data exploration applications in R, limiting the functionality to the widgets and libraries in Shiny.

OpenCPU is a system for embedded scientific computing and reproducible research.[6] Similar to the compute service in Kvik, it offers an HTTP API to the R programming language to provide an interface with statistical methods. It allows users to make function calls to any R package and retrieve the results in a wide variety of formats such as JSON or PDF. Users can chose to host their own R server or use public servers, and OpenCPU works in a single-user setting within an R session, or a multi-user setting facilitating multiple parallel requests. This makes OpenCPU suitable for building a service that can run statistical analyses. OpenCPU provides a Javascript library for interfacing with R, as well as Docker containers for easy installation. OpenCPU has been used to build multiple applications.^26^. The compute service in Kvik follows many of the design patterns in OpenCPU. Both systems interface with R packages using a hybrid state pattern over HTTP. Both systems provide the same interface to execute analyses and retrieve results. While OpenCPU is implemented on top of R and Apache, Kvik is implemented from the ground up in Go. Because of the similarities in the interface to R in Kvik we provide packages for interfacing with our own R server or OpenCPU R servers

Renjin is a JVM-based interpreter for the R programming language.[7] It enables integrating the R interpreter in web applications. Since it is built on top of the JVM it allows developers to write data exploration applications in Java that interact directly with R code, both running on top of the JVM. Although Renjin provides as a service pre-built versions of packages from CRAN and BioConductor, not all packages can be built for use in Renjin. This makes it necessary to re-implement the packages that cannot be built for use in Renjin.

### Parallel and Distributed Execution

Biogo is a bioinformatics library written in Go. It provides functionality to analyze genomic and metagenomic datasets in the Go programming language.[3] Using the Go programming language the developers are able to provide high-performance parallel processing in a clean and simple programming language. Go provides the ease of programming as popular scripting languages such as R, but the performance of low-level langauges such as C++.

MapReduce is a popular framework and programming model to analyze large-scale datasets using distributed and parallel computing.[8] While MapReduce has shown its usefulness in certain applications, it does not fit the different statistical methods in bioinformatics. An alternative to MapReduce is Spark, which supports high-performance parallel execution of iterative machine learning methods by partitioning data across a set of machines and optimizing the data access.[9] Spark is growing in popularity, especially with the Spakrlyr^27^ and SparkR^28^ R packages that allow R users to run analyses on top of Spark without having to port the analysis code to another language.

ADAM and VariantSpark are systems that are implemented on top of Spark to analyse genomic data. ADAM provides a set of formats, APIs and processing stage implementations [10] while VariantSpark provides an method for clustering genomic variants[11].

Pachyderm^29^ is a system for running containerized data analysis pipelines. Each step in an analysis pipeline is run within a software container and the output data is version controlled. Pachyderm automatically partitions and distributes the input data to enable parallel processing. Pachyderm runs on top of Kubernetes^30^, a system for managing and deploying containerized applications.

### Visualization tools

Cytoscape is an open source software platform for visualizing complex networks and integrating these with any type of attribute data[12]. It allows for analysis and visualization in the same platform. Users can add additional features, such as databases connections or new layouts, through Apps. One such app is cyREST which allows external network creation and analysis through a REST API[13]. To bring the visualization and analysis capabilities to the web the creators of Cytoscape have developed Cytoscape.js^31^, a Javascript library to create interactive graph visualizations.

Caleydo is a framework for building applications for visualizing and exploring biomolecular data[?]. Until 2014 it was a standalone tool that needed to be downloaded, but the Caleydo team are now making the tools web-based. There have been several applications built using Caleydo: StratomeX for exploring stratified heterogeneous datasets for disease subtype analysis[14]; Pathfinder for exploring paths in large multivariate graphs[15]; UpSet to visualize and analyse sets, their intersections and aggregates[16]; Entourage and enRoute to explore and visualize biological pathways [17][18]; LineUp to explore rankings of items based on a set of attributes[19]; and Domino for exploring subsets across multiple tabular datasets[20].

BioJS is an open-source JavaScript framework for biological data visualization.[21] It provides a community-driven online repository with a wide range components for visualizing biological data contributed by the bioinformatics community. BioJS builds on node.js^32^ providing both server-side and client-side libraries.

### Containerized analysis

In the later years software containers have been widely adopted by both the software industry as well as research communities. Containers provide an isolated execution environment that can be used to package and run an application with all its dependencies, library versions and configuration files. In researchers containers have become popular because they provides a reproducible environment that can be shared between research projects.

In bioinformatics researchers are starting to bundle their software using software containers, such as Docker. There are repositories such as BioContainers[22] and BioBoxes[23] that provide containers preinstalled with software for doing analyses and running different applications. Systems such as Galaxy now allow researchers to build analysis pipelines where each step is executed within a software container.

### Kvik and Kvik Pathwys

We have previously built a system for interactively exploring gene expression data in context of biological pathways.[24] Kvik Pathways is a web application that integrates gene expression data from the Norwegian Women and Cancer (NOWAC) cohort together with pathway images from the Kyoto Encyclopedia of Genes and Genomes (KEGG). We used the experience building Kvik Pathways to completely re-design and re-implement the R interface in Kvik. From having an R server that can run a set of functions from an R script, it now has a clean interface to call any function from any R package, not just retrieving data as a text string but in a wide range of formats. We also rebuilt the database interface, which is now a seperate service. This makes it possible to leverage its caching capabilities to improve latency. This transformed the application from being a single monolithic application into a system that consists of a web application for visualizing biological pathways, a database service to retrieve pathway images and other metadata, and a compute service for interfacing with the gene expression data in the NOWAC cohort. We could then re-use the database and the compute service in the MIxT application.

## Discussion

We argue that developing data exploration applications using a microservice architecture is a viable alternative to the traditional monolithic approach. By packaging each component of an application in a software container application developers can reuse and share parts of an application across research teams and projects.

Although the partition of components can help break up an application into manageable parts, there is more overhead with deploying and monitoring these than a single application. Leslie Lamport’s famous quote *You know you have a distributed system when the crash of a computer you’ve never heard of stops you from getting any work done*. perfectly describes the possible challenges application developers face when moving into a microservice architecture. Monitoring the health of the different services and keeping the services running is a challenge, but systems such as Kubernetes^33^ provide the necessary functionality to manage containerized applications.

### Future work

Although we have a first working prototype of the microservices and the MIxT web application, there are a few points we aim to address in future work.

The first issue is to improve the user experience in the MIxT web application. Since it is performing many of the analyses on demand, the user interface may seem unresponsive. We are working on mechanisms that gives the user feedback when the computations are taking a long time.

The database service provides a sufficient interface for the MIxT web application. While we have developed the software packages for interfacing with more databases, these haven’t been included in the database service yet. In future versions we aim to make the database service be a interface for all our applications. We also aim to improve how we capture data provenance. We aim to provide database versions and meta-data about when a specific item was retrieved from the database.

One large concern that we haven’t addressed in this paper is security. In particular one security concern that we plan to address in Kvik is the restrictions on the execution of code in the compute service. We plan to address this in the next version of the compute service, using methods such as AppArmor^34^ that can restrict a program’s resource access.

We also aim to explore different avenues for scaling up the compute service. Since we already interface with R we can use the Sparklyr or SparkR packages to run analyses on top of Spark. Using Spark as an execution engine for data anlyses will enable applications to explore even larger datasets.

## Conclusions

We have designed an approach for building data exploration applications in systems biology that builds on a microservice architecture. Using this approach we have built a web application that leverages this architecture to integrate statistical analyses, interactive visualizations, and data from biological databases. While we have used our approach to build an application in systems biology, we believe that the microservice architecture can be used to build data exploration systems in other disciplines as well.

## Footnotes

1 software.broadinstitute.org/gsea/msigdb

2 kegg.jP

3 cran.r-project.org

4 bioconductor.org

5 galaxyproject.org

6 commonwl.org

7 docker.com

8 coreos.com/rkt

9 biocontainers.pro

10 github.com/vdumeaux/mixtR

11 aws.amazon.com

12 eutils.ncbi.nlm.nih.gov

13 software.broadinstitute.org/gsea/msigdb

14 genenames.org

15 kegg.jp

16 github.com/vdumeaux/mixtR

17 github.com/fjukstad/gocache

18 Available at hub.docker.com/r/fjukstad/kvik-r and hub.docker.com/r/fjukstad/db

19 sigmajs.org

20 cran.r-project.org/web/packages/GGally

21 d3js.org

22 See aws.amazon.com/ec2/instance-types for more information about AWS EC2 instance types.

23 shiny.rstudio.com

24 leafletjs.com

25 threejs.org

26 opencpu.org/apps.html

27 spark.rstudio.com

28 spark.apache.org/docs/latest/sparkr.html

29 pachyderm.io

30 kubernetes.io

31 js.cytoscapejs.org

32 nodejs.org

33 kubernetes.io

34 wiki.ubuntu.com/AppArmor

## References

[1] A. Sboner, X. J. Mu, D. Greenbaum, R. K. Auerbach, and M. B. Gerstein, “The real cost of sequencing: higher than you think!,” Genome biology, vol. 12, no. 8, p. 125, 2011.

[2] P. J. Cock, T. Antao, J. T. Chang, B. A. Chapman, C. J. Cox, A. Dalke, I. Friedberg, T. Hamelryck, F. Kauff, B. Wilczynski, et al., “Biopython: freely available python tools for computational molecular biology and bioinformatics,” Bioinformatics, vol. 25, no. 11, pp. 1422–1423, 2009.

[3] R. D. Kortschak and D. L. Adelson, “b?ogo: a simple high-performance bioinformatics toolkit for the go language,” bioRxiv, 2014.

[4] C. L. Williams, J. C. Sica, R. T. Killen, and U. G. Balis, “The growing need for microservices in bioinformatics,” Journal of Pathology Informatics, vol. 7, 2016.

[5] V. Dumeaux, B. Fjukstad, H. Fjosne E J.-O. Frantzen, M. Muri Holmen, E. Rodegerdts, E. Schlichting, A.-L. Børresen-Dale, L. A. Bongo, E. Lund, and M. T. Hallett, “Interactions between the tumor and the blood systemic response of breast cancer patients,” Under review, 2017.

[6] J. Ooms, “The opencpu system: Towards a universal interface for scientific computing through separation of concerns,” arXiv preprint arXiv:1406.4806, 2014.

[7] A. Bertram, “Renjin: The new r interpreter built on the jvm,” in The R User Conference, useR! 2013 July 10-12 2013 University of Castilla-La Mancha, Albacete, Spain, vol. 10, p. 105, 2013.

[8] J. Dean and S. Ghemawat, “Mapreduce: simplified data processing on large clusters,” Communications of the ACM, vol. 51, no. 1, pp. 107–113, 2008.

[9] M. Zaharia, M. Chowdhury, M. J. Franklin, S. Shenker, and I. Stoica, “Spark: Cluster computing with working sets.,” HotCloud, vol. 10, no. 10–10, p. 95, 2010.

[10] M. Massie, F. Nothaft, C. Hartl, C. Kozanitis, A. Schumacher, A. D. Joseph, and D. A. Patterson, “Adam: Genomics formats and processing patterns for cloud scale computing,” University of California, Berkeley Technical Report, No. UCB/EECS-2013, vol. 207, 2013.

[11] A. R. O’Brien, N. F. Saunders, Y. Guo, F. A. Buske, R. J. Scott, and D. C. Bauer, “Variantspark: population scale clustering of genotype information,” BMC genomics, vol. 16, no. 1, p. 1052, 2015.

[12] P. Shannon, A. Markiel, O. Ozier, N. S. Baliga, J. T. Wang, D. Ramage, N. Amin, B. Schwikowski, and T. Ideker, “Cytoscape: a software environment for integrated models of biomolecular interaction networks,” Genome research, vol. 13, no. 11, pp. 2498–2504, 2003.

[13] K. Ono, T. Muetze, G. Kolishovski, P. Shannon, and B. Demchak, “Cyrest: Turbocharging cytoscape access for external tools via a restful api,” F1000Research, vol. 4, 2015.

[14] A. Lex, M. Streit, H.-J. Schulz, C. Partl, D. Schmalstieg, P. J. Park, and N. Gehlenborg, “Stratomex: Visual analysis of large-scale heterogeneous genomics data for cancer subtype characterization,” in Computer graphics forum, vol. 31, pp. 1175–1184, Wiley Online Library, 2012.

[15] C. Partl, S. Gratzl, M. Streit, A. M. Wassermann, H. Pfister, D. Schmalstieg, and A. Lex, “Pathfinder: Visual analysis of paths in graphs,” in Computer Graphics Forum, vol. 35, pp. 71–80, Wiley Online Library, 2016.

[16] A. Lex, N. Gehlenborg, H. Strobelt, R. Vuillemot, and H. Pfister, “Upset: visualization of intersecting sets,” IEEE transactions on visualization and computer graphics, vol. 20, no. 12, pp. 1983–1992, 2014.

[17] A. Lex, C. Partl, D. Kalkofen, M. Streit, S. Gratzl, A. M. Wassermann, D. Schmalstieg, and H. Pfister, “Entourage: Visualizing relationships between biological pathways using contextual subsets,” IEEE transactions on visualization and computer graphics, vol. 19, no. 12, pp. 2536–2545, 2013.

[18] C. Partl, A. Lex, M. Streit, D. Kalkofen, K. Kashofer, and D. Schmalstieg, “enroute: Dynamic path extraction from biological pathway maps for in-depth experimental data analysis,” in Biological Data Visualization (BioVis), 2012 IEEE Symposium on, pp. 107–114, IEEE, 2012.

[19] S. Gratzl, A. Lex, N. Gehlenborg, H. Pfister, and M. Streit, “Lineup: Visual analysis of multiattribute rankings,” IEEE transactions on visualization and computer graphics, vol. 19, no. 12, pp. 2277–2286, 2013.

[20] S. Gratzl, N. Gehlenborg, A. Lex, H. Pfister, and M. Streit, “Domino: Extracting, comparing, and manipulating subsets across multiple tabular datasets,” IEEE transactions on visualization and computer graphics, vol. 20, no. 12, pp. 2023–2032, 2014.

[21] J. Gómez, L. J. García, G. A. Salazar, J. Villaveces, S. Gore, A. García, M. J. Martín, G. Launay, R. Alcantara, N. D. T. Ayllón, et al., “Biojs: an open source javascript framework for biological data visualization,” Bioinformatics, p. btt100, 2013.

[22] F. da Veiga Leprevost, B. A. Grüning, S. A. Aflitos, H. L. Röst, J. Uszkoreit, H. Barsnes, M. Vaudel11, P. Moreno, L. Gatto13, J. Weber, et al., “Biocontainers: An open-source and community-driven framework for software standardization,”

[23] P. Belmann, J. Dröge, A. Bremges, A. C. McHardy, A. Sczyrba, and M. D. Barton, “Bioboxes: standardised containers for interchangeable bioinformatics software,” GigaScience, vol. 4, no. 1, p. 47, 2015.

[24] B. Fjukstad, K. S. Olsen, M. Jareid, E. Lund, and L. A. Bongo, “Kvik: three-tier data exploration tools for flexible analysis of genomic data in epidemiological studies,” F1000Research, vol. 4, 2015.

